# Breedbase: An Integrated Breeding Informatics System with Statistical Analysis and Decision-Support Tools

**DOI:** 10.64898/2026.11.11.765829

**Authors:** Isaak Y Tecle, Titima Tantikanjana, Christiano Simões, Clay Birkett, Amanda Hulse-Kemp, Jean-Luc Jannink, Srikanth Kumar Karaikal, Keo Corak, Amber Lockrow, Heather Manching, Ben Maza, Naama Menda, Christine Nyaga, Prasad Peteti, Adrian Powell, Ryan Preble, Ismail Y Rabbi, Mercedes Thieme, David Waring, Wolfgang Zierer, Lukas A. Mueller

## Abstract

Breeding programs are becoming increasingly data-intensive, digital and collaborative. One-stop digital breeding ecosystems, in which data storage, field workflow tools, statistical analysis and breeding decision-support tools are seamlessly integrated, are increasingly important. Breedbase (https://breedbase.org) is one of the few open-source platforms that integrate these functionalities within a single system. It provides tools for phenotypic data quality control, descriptive, inferential and predictive statistical analyses. Integrated into the database are tools for single-trial analysis of variance (ANOVA), linear mixed modeling, stability analysis, correlation analysis, principal component analysis, k-means and hierarchical clustering, kinship and inbreeding analysis, genome-wide association study (GWAS), and genomic prediction. Input data and outputs from these analysis tools are visualized in interactive plots and are also available in text files for downloading and sharing. By persisting analysis datasets in the database and applying standardized and automated data preprocessing and analysis workflows, Breedbase is designed to improve efficiency, help maintain data integrity, and support reproducible analyses that can be readily shared among collaborators. Breedbase is open-source software, containerized using Docker, and available for self-hosting or cloud hosting and management by the Mueller lab.

## Background

Breeders are becoming increasingly reliant on databases and web-based applications to manage data and make evidence-driven decisions in their data-intensive breeding programs [1]. Breeding is also becoming a collaborative endeavor involving researchers from diverse time zones who need a central platform to share information and analysis results. A number of public databases are meeting breeders’ data management needs [2–5]. However, most of them are limited to only warehousing data or limited in scope in their statistical analyses, which are mostly genomic analyses [6, 7]. Therefore breeders generally download or import their datasets from the databases and use external software for analysis [8]. This separation of information warehousing and analysis is inefficient and error prone and could lead to data corruption [9]. A database with integrated data management and analysis tools would improve data integrity, reproducibility of analysis results, and efficiency. Additionally, it eases sharing of analysis input and output among collaborators.

Breedbase (https://breedbase.org) is a comprehensive database capable of storing phenotypic and genomics data [10], high-dimensional information such as spectral images and near-infrared spectroscopy (NIRS), and other breeding experiment records [10, 11]. It stores germplasm data, ontologies, phenotype and genotype data. It is also widely used for fieldwork activities, including experiment design, barcoding, treatments, and interaction with trial phenotype data collection tools such as the Field Book app [12]. The database has an intuitive web interface for querying stored information and to make use of a suite of breeding workflow tools. In addition, Breedbase supports a standardized data exchange interface called Breeding Application Programming Interface (BrAPI) [13].

Breedbase [10] is used by an increasing number of public and commercial breeding programs. Some of the projects powered by Breedbase are the Root, Tuber, and Banana (RTB) crops breeding programs (https://cassavabase.org,https://sweetpotatobase.org, https://musabase.org), Cassava Source-Sink project (https://cassabase.org), USDA Triticeae Coordinated Agricultural Project (CAP) (https://triticeaetoolbox.org/), and the Solanaceae Genomics Network (SGN, https://solgenomics.net) [14]. There are also an increasing number of private seed companies and academic research labs that use the database in private networks.

The ultimate goal of Breedbase is to provide a one-stop breeding informatics ecosystem where users can perform the activities mentioned above and elaborated previously by [10], and tools for statistical analysis and breeding decision support. Towards this end, Breedbase has been developing and integrating a variety of statistical tools that can be used to analyze the information stored in its database. A few of the analysis tools have been covered in previous publications [11, 15, 16]. The tools implement statistical analyses ranging from basic data exploration to inferential statistics to predictive statistics. The objective of this paper is to document new statistical analysis tools implemented in Breedbase and describe upgrades to previously published ones. To demonstrate the tools in this paper CassavaBase (https://cassavabase.org), the NextGen Cassava Breeding Project (https://nextgencassava.org) database, and a flagship Breedbase instance, will be used. All URLs to the analysis tools will point to CassavaBase, but all the tools described here are available in any Breedbase instance.

## Implementation

### Software

The software dependencies of Breedbase are documented in our previous publication [10]. Here we focus on the software used for data analysis and visualization. The backbone of most data analysis is implemented using the R language [17], in which standard, contributed, and in-house developed packages are used. Data pre-processing and static plot generation are done using a collection of R packages called tidyverse (dplyr, tidyr, ggplot2 (https://tidyverse.org)) [18]; its extensions, such as ggfortify [19], ggtree [20] and treeio; and data.table [21]. In-house developed R packages, phenoAnalysis and genoDataFilter (https://github.com/solgenomics/rPackages), are also used for the phenotype and genotype data pre-processing. Other analysis-specific R packages are lme4 [22], sommer [23], lmerTest [24], lsmeans/emmeans [25], and agricolae [26]. Interactive data visualizations are generated using D3 (https://d3js.org).

To ensure long-term maintainability of the Breedbase system–including the analysis tools–automated tests, such as unit and integration tests, are built into the code base. At the time of writing, the automated test suite comprises approximately 10,000 test assertions, which are updated alongside code changes and run on every pull request before it is merged into the main code base and released, helping to detect errors and regressions early.

To facilitate the use of Breedbase tools, including the analysis tools, an online user manual (https://solgenomics.github.io/sgn/) is continuously updated.

### Statistics

Following data upload to a BreedBase instance, users can perform descriptive, inferential, and predictive statistical analyses. Intuitive web-based graphical user interfaces (GUIs) enable users to query, organize, and structure datasets and to conduct analyses using point-and-click options. Results are presented through interactive visualizations. Users can download outputs, text files, and plots and, in certain cases, save them to the database for subsequent downstream analysis or retrieval.

Below, we provide an overview of the various statistical analysis tools available in Breedbase.

### Data Preparation for Analysis

Depending on the analysis type, users can perform analysis on data from a single trial or a combination of trials. If the data of interest is a single trial, inclusive of all accessions and plots, users can simply search and go to the trial detail page to view the trial data and perform analysis. Alternatively, field trials can be found via the Search Wizard (e.g., https://cassavabase.org/breeders/search), where one or multiple trials can be selected fully or subsetted by certain criteria (e.g., accession names and treatments) and saved as a list or dataset. A list is a one-dimensional data structure that could be a list of trials, traits, or locations. On the other hand, a dataset can be created by combining different dimensions of data—such as breeding programs, genotyping protocols, trials, locations, years, accessions, traits, and plots—among others [10]. In addition, users can compose their datasets with subsets of accessions or plots from the trials included.

Once a dataset or a list is created, it becomes available for analysis on any analysis tool page provided that the user is logged in to the database. Users can go to an analysis tool homepage, select their dataset, and perform phenotype data quality controls and data analysis. Lists are only available in some analysis tools. The advantage of organizing data in lists and datasets is that they persist in the database. Furthermore, they can be set private (viewable by the owner only) or public (viewable by any logged in user, a helpful option for sharing data with collaborators) at any time.

For this manuscript and to demonstrate the analysis tools, a dataset called BIOINFOR-IYT-26 (https://cassavabase.org/dataset/366) was created. The dataset consists of 796 cassava accessions from two trials, phenotyped for 33 traits at two locations and genotyped with 107,825 SNPs. A subset from the BIOINFOR-IYT-26 dataset, prepared for kinship and hierarchical clustering analysis, is also available as BIOINFOR-IYT-26-KINSHIP (https://cassavabase.org/dataset/365). All analysis outputs shown in the figures (Fig 1–6) were generated using these datasets.

**Figure 1:**
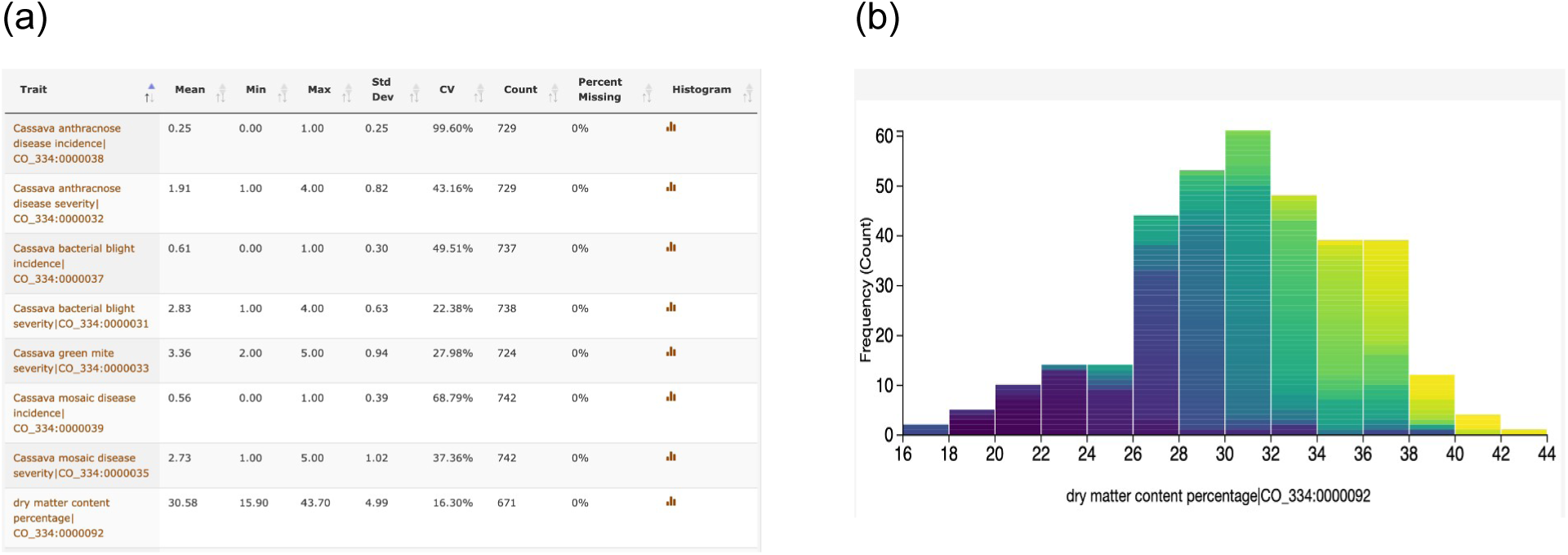
(a) Descriptive statistics for each phenotyped trait in a trial. (b) An example visualization of the distribution of trait data in an interactive histogram.

**Figure 2:**
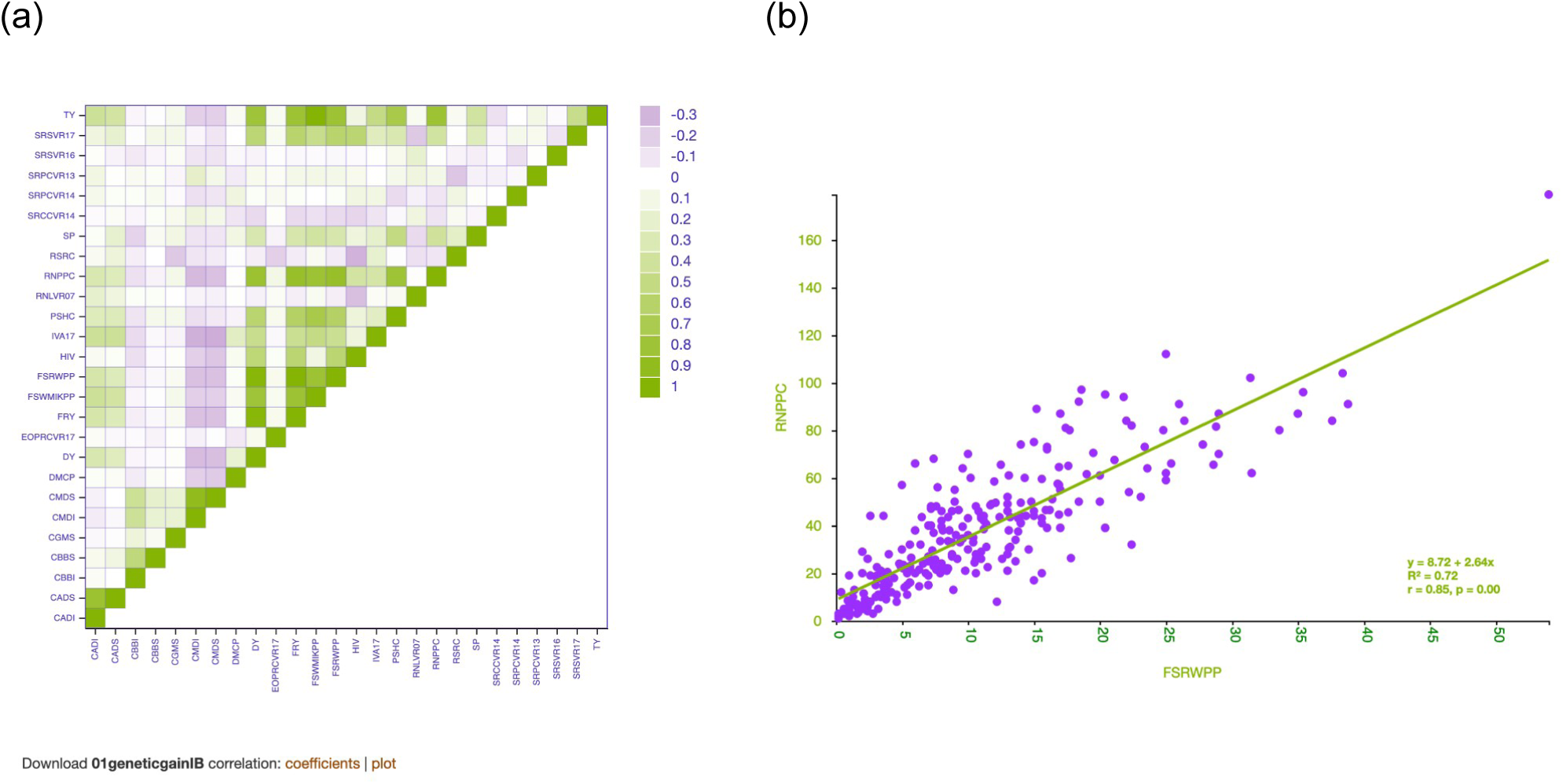
(a) An interactive heatmap from Pearson correlation analysis. Pointing the cursor at a cell representing a pair of traits shows the correlation coefficient and p-value. (b) Scatter plot of data for a pair of traits data with an ordinary least squares regression line.

**Figure 3:**
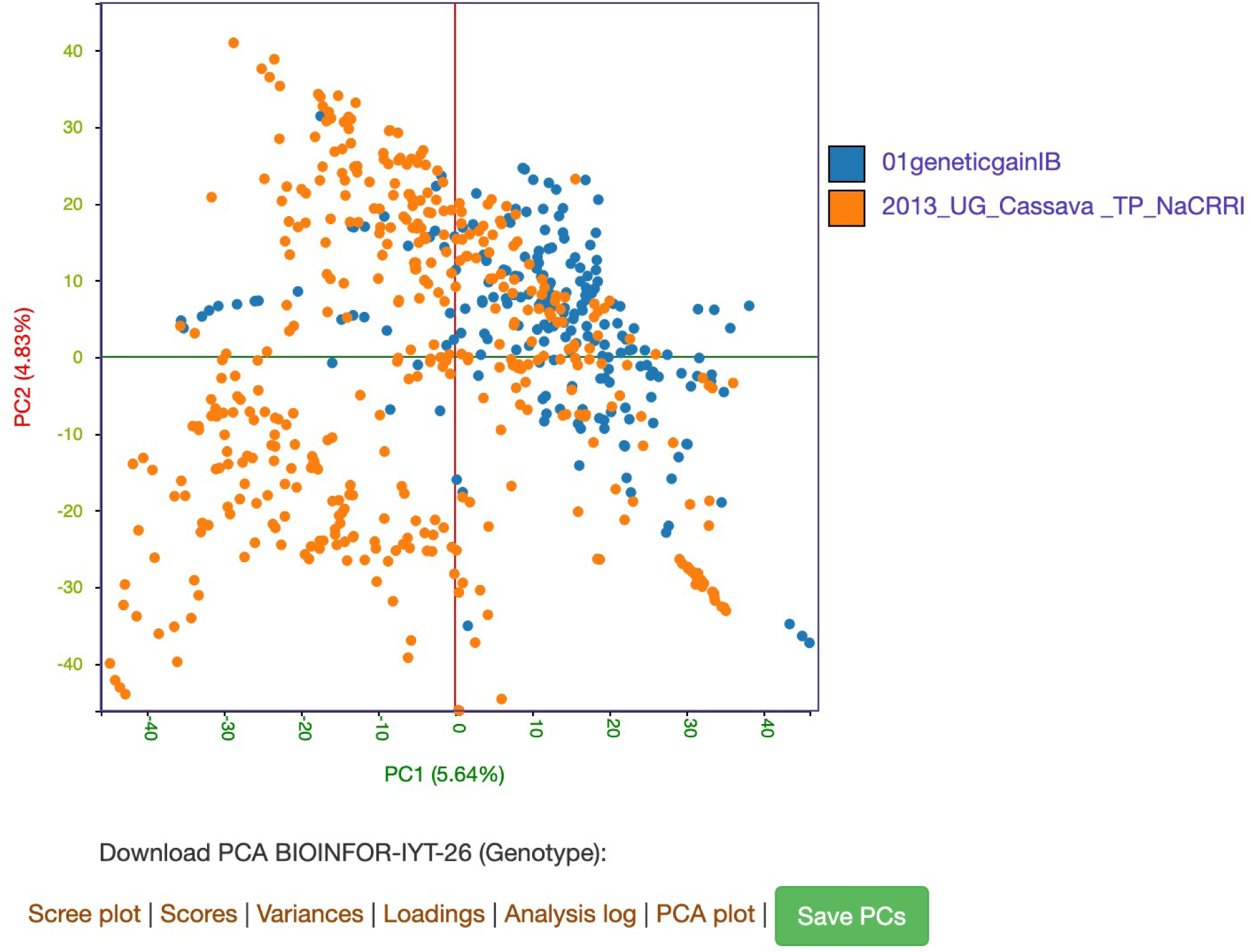
An interactive plot depicting scores from a principal component analysis (PCA). Scores are color-coded when individuals belong to different categories, such as trials. Links are provided for downloading the complete PCA output, including Scree plot. The ‘Save PCs’ link allows the top ten principal components to be saved to the database.

**Figure 4:**
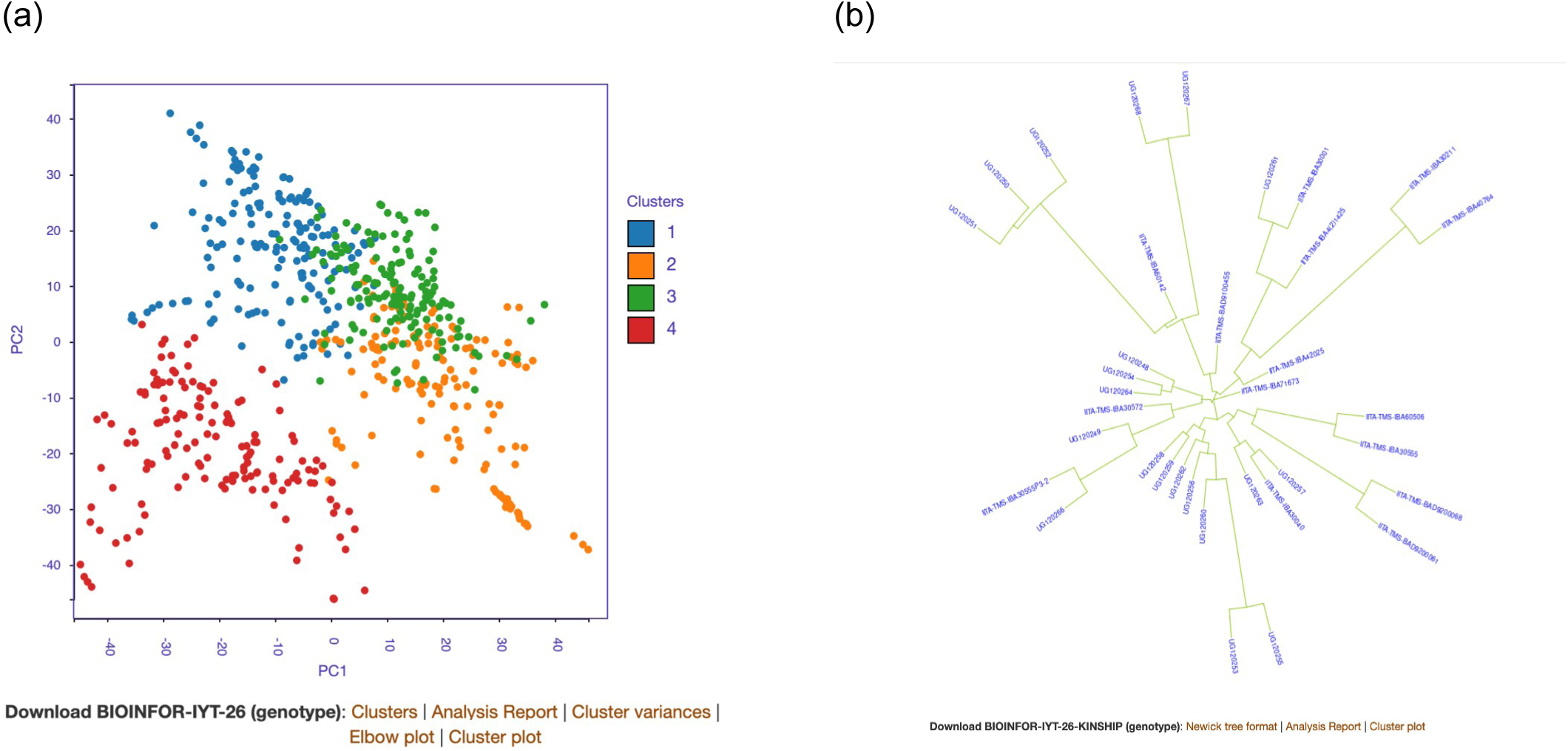
Outputs from K-means (a) and hierarchical (b) clustering analyses. In addition to the plots, links are provided to download the analysis output, diagnostic plot, and analysis report.

**Figure 5:**
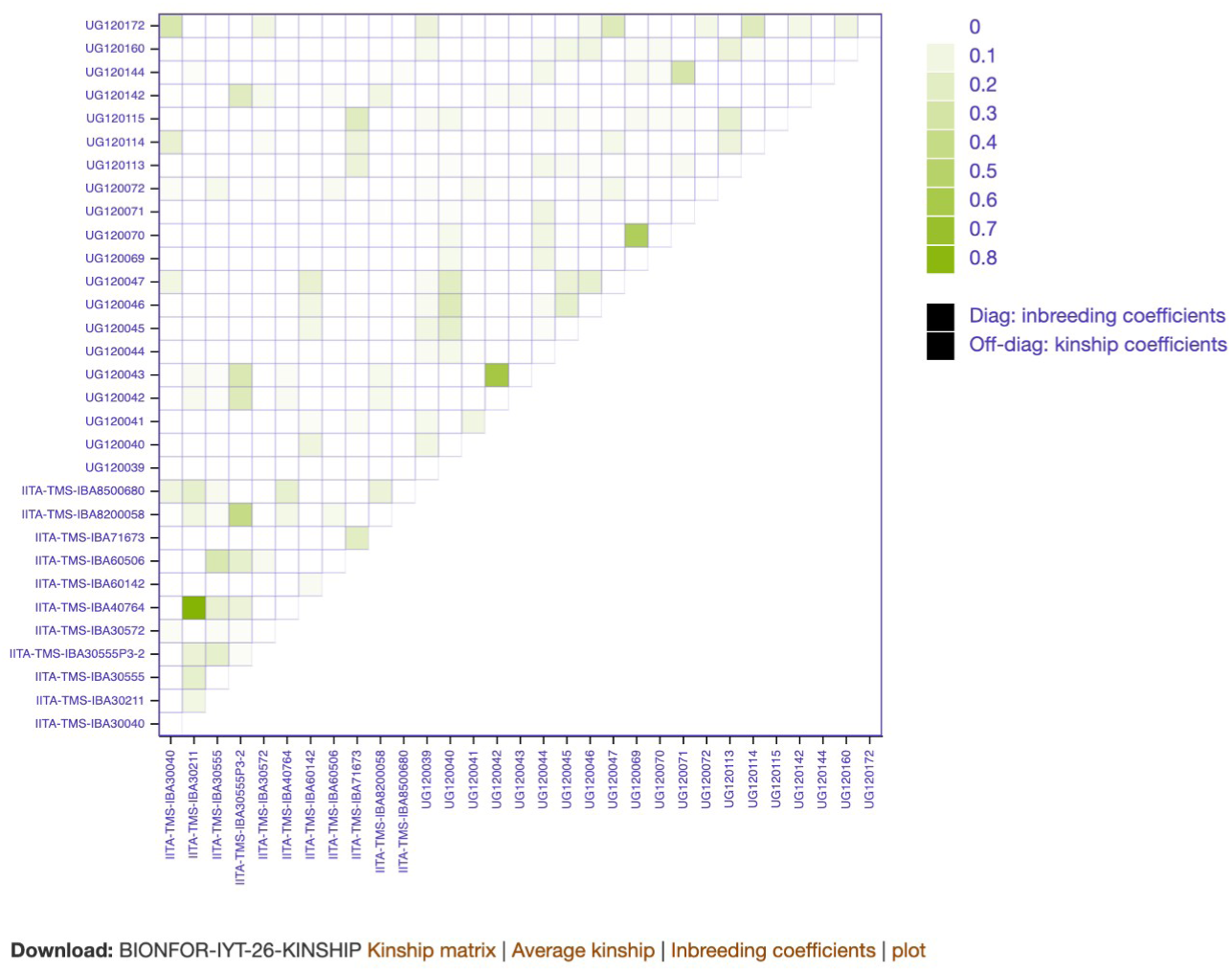
An interactive heatmap visualizing pairwise kinship off-diagonal cells and inbreeding coefficient of each clone in the diagonal cells.

**Figure 6.**
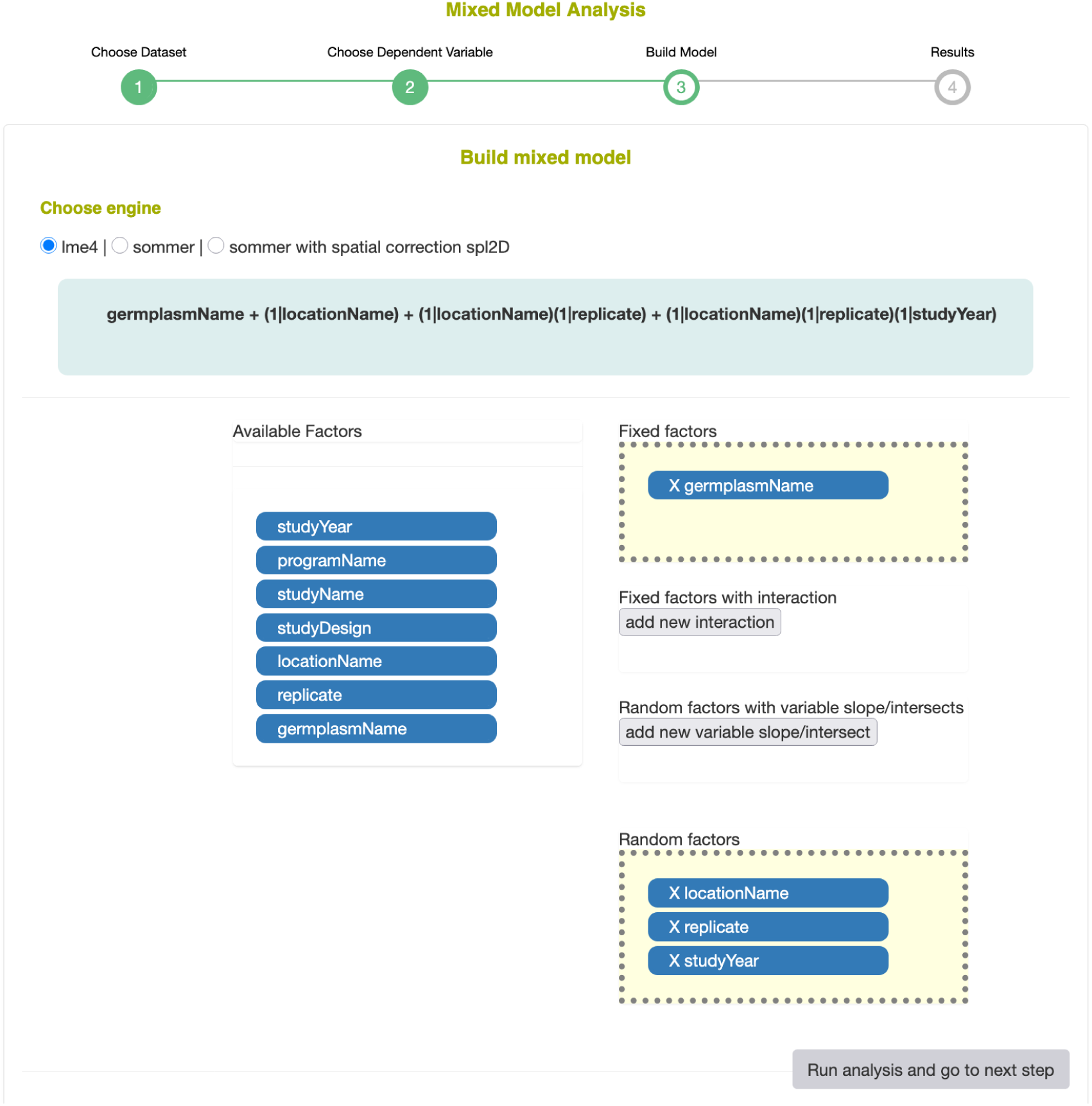
Mixed model interface for selecting the modeling engine and specifying fixed effects, random effects, and interaction terms by drag and drop, with the resulting model formula displayed on the page.

### Availability of Analysis Tools

Analysis and decision-support tools in Breedbase can be accessed through several entry points. Most tools, including correlation, clustering, kinship, genomic prediction (solGS), and stability analysis, are available from the Analyze menu of a Breedbase instance. Some analyses are also accessible directly from trial detail pages, including ANOVA, correlation analysis, and principal component analysis (PCA). In addition, selected tools are embedded within analysis and decision-support workflows.

For example, ANOVA, correlation analysis, PCA, kinship analysis, and clustering are available within the solGS pipeline, while PCA and kinship analysis can be accessed as part of the solGWAS workflow. The tools displayed in the Analyze menu may vary among Breedbase instances depending on their configuration; tools that are not displayed can be enabled when required.

## Results

### Exploratory Data Analysis

#### Descriptive statistics

Descriptive statistics can be accessed in various places depending on how the data is structured. If the user is interested in a single trial only, they can go to the trial’s detail page, scroll to the ‘Phenotypic Summary Statistics’ section, and view the descriptive statistics for the trial data. This includes the mean, minimum and maximum, standard deviation, coefficient of variation, and percentage of missing values for each trait phenotyped in the trial (Fig 1a). When the data is structured as a dataset or list, the summary data is shown in the pipeline of the analysis where the data is used as an input.

The distribution of trait data can be viewed in an interactive histogram, which displays the observation units in each data bin (Fig 1b), or boxplots as described more in the phenotype quality control section below. If users want to filter and store observation units within a selected trait values range, they can use a tool called ‘Graphical Filtering’ (https://cassavabase.org/tools/graphicalfiltering), which currently works for trials organized as a list. The filtering of observation units can be done based on a selected data range of a single trait or combinations of traits. Additionally, if users wish to determine outliers and mask them in downstream analysis, they can use a ‘Quality Control (QC)’ tool (https://cassavabase.org/tools/qualitycontrol), as described by [26], which enables them to view trait data distribution using a boxplot and filter out outliers using the interquartile range method. Users can use the same tool also to compare distribution of trait data across multiple trials.

To estimate relationships between traits, users can run Pearson correlation analyses, based on the `rcor.test()` function (method= pearson, use=pairwise, on accession means) [27], which generates an interactive heatmap depicting the relationship magnitude, direction, and statistical significance (p-value) (Fig 2a). They can point their cursor at a cell representing a pair of traits and view the correlation coefficient and p-value. Additionally, a scatter plot of the paired traits data is rendered with an ordinary least squares regression line, which can reveal more insights on the bivariate relationship (Fig 2b).

#### Principal Component Analysis (PCA)

In addition to low-dimensional phenotypic data, Breedbase is optimized to store high-dimensional data, including Near-Infrared Spectroscopy (NIRS) [11], transcriptomic, metabolomic, and marker data. Using high-dimensional data in statistical models to drive breeding decisions may be computationally intensive and time consuming. Importantly, high-dimensional data may be noisy and contain variables with high multicollinearity or spurious correlation and is difficult to visualize [28]. Due to potential multicollinearity among variables, identifying patterns and causal factors becomes unreliable. A common approach is to reduce the high-dimensional data into a manageable number of representative variables, while maintaining most of the variance in the original data, before modeling or any statistical analysis. Principal component analysis (PCA), an unsupervised machine learning technique, is one of the most popular approaches for this purpose [29].

Additionally, PCA can uncover underlying population structure in a dataset. The PCA loadings can be used to estimate the direction and magnitude of the effects of the original variables on principal components. The principal component loadings, in combination with the scores, can be used to explain the subgrouping of the individuals in a dataset and identify the variables underlying it [30].

The PCA tool implements the `prcomp()` function from the base R stats package. Prior to running the PCA, data is cleaned for missing data, filtered for variables with constant variance, averaged in case of phenotype data, scaled, and centered. The data processing is logged in a file that users can download.

The PCA output includes an interactive plot of the top two principal components (PCs) in the browser, allowing users to identify individual data points. For datasets composed of individuals from multiple trials, data points are color coded according to their trial of origin. Additionally, the PCA tool generates downloadable static PCs, scree plots, scores for the top ten PCs, eigenvalues (representing the variance explained by the PCs), and loadings (representing the direction and magnitude of the relationships between the variables and the PCs) (Fig 3). Users also have the option to store the top ten PCs in the database for use in downstream statistical analyses or for later retrieval.

#### Clustering

Efficient crossing designs and genetic gain depend on reliably partitioning breeding materials into genetically distinct groups and crossing individuals from genetically different groups [31]. This becomes challenging when individuals are phenotyped on multiple traits or assayed across other high-dimensional variables, such as markers or metabolites. Clustering algorithms are helpful in partitioning the dataset individuals into a desired number of groups (k), e.g., using k-means clustering; alternatively, hierarchical clustering can visualize the clustering of individuals in a tree-like structure. Breedbase implements both k-means and hierarchical clustering. Users can use the web form to select a dataset, specify the data type (phenotype or genotype) for clustering, choose either the k-means or hierarchical method, and in the case of kmeans, fill in the number of clusters (k) into which the dataset should be partitioned. Users can use both clustering methods on datasets with quantitative phenotype data and genetic marker data. When using the genomic prediction tool (solGS), described in a later section, users can perform cluster analysis based on genomic estimated breeding values (GEBVs) of multiple traits. If users calculate selection indices using the selection index calculator integrated in the genomic prediction pipeline, they can also select a fraction of the indexed individuals, depending on their choice of selection intensity, and perform clustering on that subset using the marker data of the selected fraction.

Prior to applying a clustering method, data is filtered for missing values, minor allele frequency (MAF, >0.01), and monomorphic markers in case of genetic marker data. Markers and individuals with greater than 20% missing values are removed. For marker-based clustering, a PCA is performed first and the top 10 PCs are used for clustering. Several R packages are used to implement the clustering algorithm. The data preprocessing is done using custom Breedbase R packages: phenoAnalysis and genoDataFilter. K-means clustering is done using the `kmeans()` function from the stats R package, with 10 times random initializations (nstart=10). For reference, the k-means clustering method also computes the recommended cluster numbers for the dataset, based on the `kmeansruns(),(runs=10),` function from the fpc R package [32]. Additionally, an elbow plot is generated using the `fviz_nbclust()` function from the factoextra R package [33], so users can judge the appropriate cluster numbers for their dataset based on within-cluster sum of squares (WSS). Hierarchical clustering is done using the standard R functions `dist()` (Euclidean method) and `hclust()` (complete method), both from the base R stats package, while the tree visualization is done using `ggtree()` function from the ggtree R package [20]. Newick tree format is generated using `as.phylo()` function from the R package ape [34].

The output of the k-means clustering method generates an interactive plot, with color-coded clusters, in the browser, cluster assignments, elbow plot, and text files of the clusters, as well as variances (within-cluster sum of squares, between-cluster sum of squares), and an analysis log can be downloaded (Fig 4a). The hierarchical clustering tool generates a static dendrogram plot, a Newick tree format representation, and an analysis log (Fig 4b). All are available for download.

#### Kinship and Inbreeding

A kinship matrix summarizes the degree of shared ancestry between pairs of individuals, while inbreeding coefficients measure the probability that the two alleles at a locus within an individual are identical by descent (IBD). Kinship matrices are commonly used as an input in statistical and breeding modeling, such as in genomic prediction and breeding value estimation, genome-wide association modeling, conventional crossing designs, and genomic prediction of cross-performance [15, 35, 36].

Kinship and inbreeding coefficients are computed using the `A.mat()` function from the rrBLUP R package [37], whereas the interactive visualization is rendered using D3 JavaScript library. Users can estimate kinship and inbreeding coefficients using genomic SNP marker data. The tool generates an interactive heat map of the kinship matrix (off-diagonal) and inbreeding coefficients (diagonal) plot on the browser, downloadable plot, text files of the kinship matrix, inbreeding coefficients, and average kinship (Fig 5).

### Inferential statistics

#### Analysis of Variance (ANOVA)

Breedbase implements analysis of variance (ANOVA) tools, which are used to compare means of two or more groups, at different levels: single-trial level (single-trial analysis) and combination of trials (multi-trial analysis including genotype by environment interactions). For single-trial analysis, users can visit the trial detail page, under ‘Analysis tools’/’ANOVA’ and select a trait of interest.The tool fits a linear mixed model according to the field trial design stored in the database, using the lmerTest R package with restricted maximum likelihood (REML). Currently supported designs are the completely randomized design (CRD), randomized complete block design (RCBD), augmented design, and alpha lattice design. Genotypes are fitted as fixed effects, while replicates and blocks are fitted as random effects, with blocks nested within replicates for the alpha lattice design. The ANOVA table reports Type III Ftests with Satterthwaite-approximated denominator degrees of freedom. Outputs include the ANOVA table, displayed in the browser and downloadable as a text file; a model summary describing the fitted model; genotype adjusted means (leastsquares means); and model diagnostic plots (residuals versus fitted values and a normal Q-Q plot of residuals). Users can also estimate trait heritability using a separate linear mixed model, in which genotypes are fitted as random effects. The model is fitted using REML, which can handle unbalanced trials and augmented designs. The model estimates genetic and residual variance components while accounting for the experimental design, including replications and blocks when applicable. The heritability formula is automatically adjusted based on the trial design, using the variance components estimated by REML. The tool outputs a table containing the variance components and broad-sense heritability for each trait in the trial.

For users who require greater control over model specification, a linear mixed model tool is provided (https://cassavabase.org/tools/mixedmodels) (Fig. 6). The tool is dataset-driven: users select a dataset, which may span multiple trials, locations, and years, and one or more traits for analysis. Design and environmental factors with more than one level in the dataset, such as genotype, location, year, trial, replicate, block, and row and column positions, are automatically listed as candidate factors.

Users then select a modeling engine and, through a drag-and-drop interface, assign factors as fixed or random effects and define fixed or random interaction terms. The resulting model formula is dynamically displayed on the page as the model is built. Three modeling engines are available: lme4 [22], sommer [23], and sommer with spatial correction. The spatial option extends the specified model by including row and column random effects within each location and a two-dimensional spline over the plot row and column coordinates (spl2Da) to account for spatial trends in the field. Each trait is analyzed separately. When genotypes are fitted as random effects, the tool returns best linear unbiased predictors (BLUPs) and adjusted means. When genotypes are fitted as fixed effects, it returns best linear unbiased estimates (BLUEs) and adjusted means.

Additionally, Breedbase provides a Stability Analysis Tool (https://cassavabase.org/tools/stability), which enables users to evaluate genotype performance across multiple environments. The tool requires the data to be structured as a dataset comprising trials from at least two environments and a common set of genotypes evaluated for the same traits. It supports multi-environment trial analysis by implementing both Additive Main Effects and Multiplicative Interaction (AMMI) and Genotype plus Genotype-by-Environment interaction (GGE) methods. The tool generates ANOVA tables, AMMI stability values (ASVs), and GGE biplots, which can be used to visualize genotype performance, assess stability, and support the interpretation and ranking of genotypes across environments.

In terms of algorithms, the linear modeling approaches (ANOVA and Linear Mixed Model tools) use the `lmer()` function, using REML and Satterthwaite, from lmerTest R package [24], whereas the stability analysis uses `AMMI()` and `GGE()` functions from R package agricolae [26]. Adjusted means are calculated using the `lsmeans()/emmeans()` function from the lsmeans/emmeans R packages [25].

#### Genome-Wide Association Studies

Associations between phenotypic observations and genetic variants can be analyzed using the Breedbase-implemented solGWAS tool (https://cassavabase.org/tools/solgwas). The genome-wide association study tool uses a dataset of accessions with phenotypic and genotypic data as input. Users can visualize the distribution of trait values in histograms and optionally exclude outliers from the dataset. Principal component scores and kinship matrices can also be included as covariates. The tool generates Manhattan plots and diagnostic Q-Q plots.

For association modeling, the tool implements the GWAS() method from the rrBLUP R package [37].

### Predictive Statistics

#### Genomic Prediction

Breedbase was initially developed primarily to support the data management and analysis for the NextGen Cassava project (https://nextgencassava.org) [38], with a particular focus on genomic selection. Accordingly, the web-based genomic prediction tool solGS was developed, as previously described [15]. Since its previous publication, solGS has been enhanced in terms of performance, data visualization, and interactivity, and has gained additional features, including options to store model outputs in the database.

Here, we briefly describe the improvements to the user experience and new features integrated into the solGS prediction pipeline.

Genomic prediction jobs are computationally intensive and may take considerable time to complete, making it unreliable to wait for the process to finish through a web browser because requests can time out. To address this, a job submission system was implemented to allow users to submit modeling jobs for background processing, and receive an email notification when the analysis is complete. Alternatively, users can monitor the status of submitted jobs from their profile page. Once a job is completed, users can access the analysis results through the links provided either in the notification email or on their profile page. To facilitate faster access to the output, a caching scheme has also been implemented.

To start the genomic prediction process, users can visit the tool’s homepage (https://cassavabase.org/solgs) and select an existing dataset, if one has already been created using the search wizard (https://cassavabase.org/breeders/search), or create a new dataset as briefly described in the data preparation for analysis section above and documented in detail by [10]. Alternatively, users can search for trials with phenotypic data, select the relevant genotyping protocol, and create a training population on the fly from the tool’s home page. However, users are encouraged to first create a dataset using the search wizard because the dataset is stored in the database and can be reused for future modeling, used in other analysis tools, or downloaded for use outside the database.

Regardless of how the training data are created, users can proceed to the training population detail page, which provides a summary of the training population, a list of the phenotyped traits for which prediction models can be fitted, and links to embedded tools such as ANOVA and Pearson correlation analysis. Before modeling, users who wish to identify the most informative traits can perform Pearson correlation and ANOVA analyses directly from the training population detail page and generate outputs and metrics described in the sections above.

To fit training models, users can select any number of traits simultaneously. The modeling output for each trait includes descriptive statistics of the phenotype data used for model training, including histograms showing the trait values; variance components and the estimated single nucleotide polymorphism (SNP) heritability; prediction accuracy based on a 10-fold cross-validation; GEBVs (Fig 7a) and genetic values; an ordinary least-squares (OLS) regression plot of the GEBVs against the observed phenotype values (Fig 7b); and an analysis log. All outputs are available for download in image and text format.

**Figure 7:**
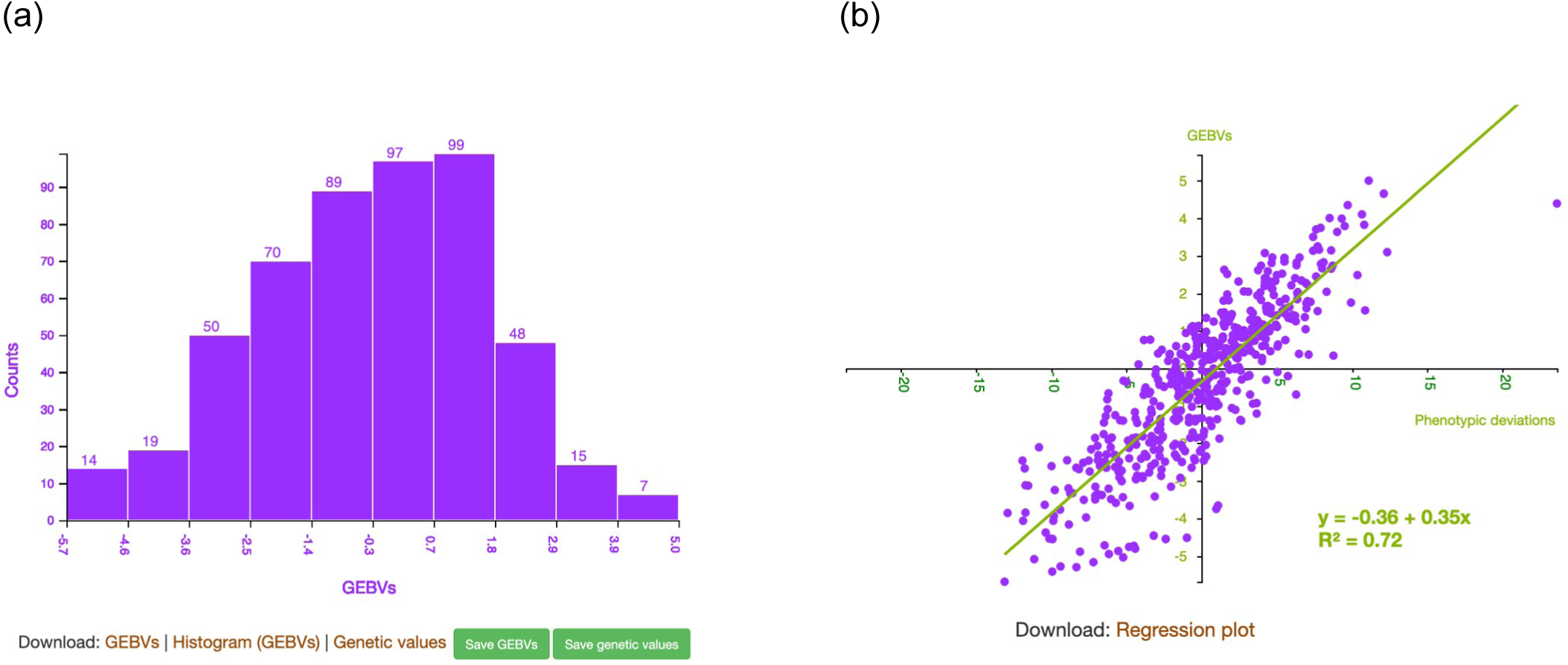
(a) Interactive histogram showing the genomic estimated breeding values (GEBVs) generated by the genomic prediction (solGS) tool. Pointing the cursor at a bar displays a table listing the individuals represented by that bar and their corresponding GEBVs. (b) Scatter plot showing the relationship between GEBVs and observed phenotypic deviations for individuals in the training population.

To predict the GEBVs of selection candidates (individuals with genotype data only), users must first identify the candidates from a section at the bottom of the model page by selecting them from a dataset or list, or by searching for a trial containing the relevant accessions, and then apply the model. The resulting GEBVs are displayed in an interactive histogram and can also be downloaded as a text file.

To support breeding decisions, the workflow integrates several diagnostic and decision-support tools. From model training through application of the trained model to predict GEBVs and genetic values of selection candidates, users can access interfaces for conducting PCA, clustering, kinship analysis, selection index analysis, genetic correlation analysis, and box plots showing expected genetic gain for the model input and output data. Users can also download the phenotypic and genotypic data used in the model, as well as the modeling results, including analysis logs. In addition, GEBVs and genetic values can be saved to the database for subsequent use.

The solGS tool builds univariate models and is implemented using the rrBLUP R package [37]. The general algorithm, including the genotypic and phenotypic data preprocessing, is described in [15]. Subsequently, the `mixed.solve()` method used for the GEBVs prediction was replaced by the `kin.blup()` method from rrBLUP version 4.6.2. An additive relationship matrix is computed from SNP marker data using the `A.mat()` function from the same package. Prediction accuracy is the mean Pearson correlation between predicted GEBVs and observed phenotypic values across 20 validation sets (10-fold cross-validation, replicated twice).

solGS includes a selection index calculator and a genetic gain estimator. The selection index calculator combines GEBVs across multiple traits into a single index by multiplying each trait’s GEBVs by a user-defined relative weight and summing the products. Because the GEBVs are not standardized and remain in their original trait units, the relative weights should account for differences in scale among traits. In addition to the index values, the calculator generates an interactive histogram of the indices and a Pearson correlation heatmap of the trait GEBVs and the indices. The genetic gain estimator compares the GEBV distributions of the training and selection populations for each trait using boxplots annotated with quartile values..

Breedbase also provides a standalone genomic prediction and cross performance (GPCP) tool [36]. The tool enables users to predict the expected performance of progeny from specific parental crosses. The method incorporates both additive and dominance effects within a mixed linear modeling framework, allowing breeders to better account for heterosis and inbreeding effects when selecting parents. This makes it especially useful in clonally propagated crops, particularly when the breeding focuses on traits where dominance plays a significant role, and breeding programs aiming to optimize crossing strategies rather than only selecting top-performing individuals.

### Links to External Analysis Services

Breedbase implements an image analysis workflow in which images can be uploaded to the database and submitted to different analysis services through a standardized image analysis API. For example, Breedbase currently interfaces with image prediction models for cassava necrosis analysis and whitefly count, developed and hosted at Makerere University, Uganda [39], as well as image-analysis pipelines developed and hosted at North Carolina State University, North Carolina, USA, including object segmentation and morphometric analysis for extracting traits such as area, perimeter, ellipse diameters, solidity, and convex hull area [40].

To use these services, users can visit the image analysis homepage of a Breedbase instance (e.g., https://cassavabase.org/tools/image_analysis),select an image stored in the database, choose an image prediction service, and submit a prediction request to the remote server. The analysis output is then returned to Breedbase, where it is stored in the database and displayed to the user. Any image analysis algorithm that conforms to the standardized application programming interface (API) can be integrated into the Breedbase workflow with minimal configuration.

### BrAPI compatibility

Breedbase is BrAPI-compliant [13], which enables interoperability with other external BrAPI-compliant tools. For example, the Field Book app [12] can connect to Breedbase through BrAPI to synchronize field layouts, traits, and collected phenotypic data between a mobile device and the database. For data analysis, the R package QBMS [41] is of particular importance, as it can conveniently access data stored in BrAPIcompliant databases directly from within an R session or script.

## Discussion

Breedbase is a one-stop breeding informatics ecosystem designed to organize, store and facilitate access to breeding data. It provides tools for breeding field workflows and integrates with external data collection applications such as the Field Book and Coordinate [42]. In addition, it includes a data analytics layer that enables breeders to analyze data directly within the database. Users can perform descriptive and inferential statistics, assess the quality of phenotypic data, including outlier detection and filtering, and perform prediction statistics such as genomic prediction. Several analysis tools that support breeding decisions are also integrated, including correlation analysis, PCA, k-means and hierarchical clustering, selection index calculation, and genetic gain estimation. Input data and analysis outputs are visualized in interactive plots and tables to facilitate interpretation and support insight generation. Both input data and analysis outputs can be downloaded, while analysis datasets and, in some cases, analysis results can be persisted in the database for subsequent use.

Breedbase distinguishes itself among publicly available databases by integrating breeding data storage, field workflow tools, data analytics, and decision-support functionality within a single platform. In contrast, many public breeding databases, such as the Enterprise Breeding System (EBS; https://ebsproject.org), Breeding Management System (BMS) [43], and DeltaBreed [4] focus primarily on data management and provide a more limited range of integrated statistical analysis tools. Stand-alone statistical analysis platforms, such as Bioflow (https://bioflow.ebsproject.org/) and Mr. Bean [8], serve a complementary role by providing analytical functionality while relying on data imported from external databases via BrAPI or uploaded from users’ local computers.

Loose coupling between data repository and analysis platforms can reduce workflow efficiency and increase the risk of data integrity issues. Integrating analysis tools with the database in which the data are stored allows data preprocessing and analysis procedures to be standardized and automated. It can also help preserve data integrity by reducing the need to transfer data files between platforms and user’s local storage systems, where errors or inconsistencies may be introduced. This integration can improve efficiency and support reproducibility of analysis results. Furthermore, integrating multiple breeding tools within one platform, as in Breedbase, enables stronger interlinking of data and facilitates seamless exploration of the related information by the user.

Most data analysis tools in Breedbase use standardized algorithms and default values for model parameters. While this facilitates consistent and streamlined analyses, it can limit users who require more nuanced or tailored analytical approaches. In the near future, we plan to introduce additional user interfaces that allow users to specify their own model parameter values, thereby increasing the flexibility of the analysis tools.

## Summary

Breedbase is a comprehensive database with a breeder data management system that seamlessly integrates tools for field workflows, statistical analysis, and breeding decision support. Integrating statistical tools directly within Breedbase can improve data analysis efficiency, reduce the risk of data integrity issues that may arise when data repository and analysis platforms are decoupled, and support reproducibility through standardized and automated workflows from data preprocessing to analysis. Input data and analysis outputs can be readily downloaded, stored in the database, and shared with collaborators. Breedbase is open-source software and individual Breedbase instances can be self-hosted or hosted in the cloud and maintained by the Mueller Lab.

## Future Directions

Although Breedbase provides an extensive suite of analytical and decision-support tools, many current methods rely on default values for model parameters. This can limit users who require more flexible or tailored analyses. Future development will therefore focus on making analytical workflows more interactive and adaptable to users’ needs by allowing users to specify their own model parameter values.

## Declarations

## Abbreviations

IITA: International Institute of Tropical Agriculture
ANOVA: Analysis of variance
GWAS: Genome-wide association study
NIRS: Near-infrared spectroscop
BrAPI: Breeding Application Programming Interface
RTB: Root, Tuber, and Banana
CAP: Coordinated Agricultural Project
SGN: Solanaceae Genomics Network
URL: Uniform Resource Locator
GUI: Graphical user interface
PCA: Principal component analysis
PC: Principal component
QC: Quality control
GEBV: Genomic estimated breeding value
MAF: Minor allele frequency
IBD: Identical by descent
REML: Restricted maximum likelihood
CRD: Completely randomized design
RCBD: Randomized complete block design
BLUP: Best linear unbiased predictor
BLUE: Best linear unbiased estimate
AMMI: Additive Main Effects and Multiplicative Interaction
GGE: Genotype plus Genotype-by-Environment interaction
SNP: Single nucleotide polymorphism
OLS: Ordinary least-squares
GCPC: Genomic prediction and cross performance
API: Application programming interface
EBS: Enterprise Breeding System
BMS: Breeding Management System

## Ethics approval and consent to participate

Not applicable.

## Consent for publication

Not applicable.

## Availability of data and materials

## Data availability

All data used to demonstrate the statistical analysis tools are stored in and publicly available through Cassavabase (https://www.cassavabase.org). The specific datasets used in this study are identified in the manuscript and are accessible to reviewers through the Cassavabase web interface. The data are made available under open-access principles in accordance with the Toronto Agreement on prepublication data sharing.

## Software Availability

The Breedbase software is open source and available on GitHub (https://github.com/solgenomics) and on Docker Hub (https://hub.docker.com/r/breedbase/breedbase). It can be used without any restrictions for commercial or non-commercial purposes. Breedbase instances can be selfhosted or hosted in the cloud and maintained by the Mueller Lab.

Project name: Breedbase

Project home page: https://breedbase.org

GitHub: https://github.com/solgenomics

Docker Hub: https://hub.docker/r/breedbase/breedbase

Operating system(s): Platform independent

Programming language: Perl, R, JavaScript, Python

License: MIT

## Authors’ contributions

IYT developed the genomic prediction, principal component analysis (PCA), clustering, kinship analysis, analysis of variance (ANOVA), and correlation analysis tools. CS developed the phenotypic data quality control, heritability and stability analysis tools. AP developed the genome-wide association study (GWAS) tool, CM developed the GPCP tool, and LAM developed the linear mixed-model tool. All other authors contributed to the development of the database and the underlying Breedbase codebase. LAM supervised the development of the statistical analysis tools. IYT drafted the manuscript. All authors contributed to discussions, reviewed the manuscript, and approved the final version.

## Acknowledgments

Breedbase has been developed through collaboration with its users and stakeholders. We thank the breeders, curators, researchers, and partner institutions whose feedback, feature requests, testing, and continued use have contributed to its development. We particularly acknowledge collaborators from IITA, CIAT, CIP, NaCRRI, NRCRI, TARI, Cornell University, the Boyce Thompson Institute, the U.S. Department of Energy Joint Genome Institute, and Makerere University.

